# A High-Precision Timing Method and Digital Interface for Closed-Loop TMS

**DOI:** 10.1101/2025.04.05.647340

**Authors:** Olli-Pekka Kahilakoski, Heikki Sinisalo, Jaakko O. Nieminen, Kyösti Alkio, Kim Valén, Gábor Kozák, Risto J. Ilmoniemi, Timo Roine

## Abstract

**Objective:** Current transcranial magnetic stimulation (TMS) protocols exhibit high inter-subject variability in treatment outcomes, highlighting the need for personalized, brain-state-dependent closed-loop stimulation protocols. To enable such protocols, we aim to provide robust, precisely timed external control of TMS, with stimulation timed relative to feedback signals such as the electroencephalogram (EEG).

**Approach:** Commercial TMS devices typically rely on trigger signals for precise external pulse timing, while adjusting stimulation parameters, such as intensity, is better handled via serial digital communication, which supports robust error detection and feedback. However, combining these communication methods is inherently complex and prone to timing issues, such as race conditions. Furthermore, trigger signals lack capabilities essential for real-time systems, such as preventing late pulse delivery.

We present a method for precise and accurate pulse timing, implemented through a digital interface that uses exclusively serial digital messaging, eliminating the need for trigger signals. This interface enables external control of pulse timing, intensity, and other parameters. The TMS device maintains its own internal clock and delivers pulses at pre-scheduled times, decoupling timing precision from the control device. Additionally, we propose a method for synchronizing such time-tracking TMS devices with commercial EEG systems, enabling precisely timed EEG–TMS.

**Main results:** Using these methods, our custom TMS device delivered pulses precisely aligned to the EEG signal, with timing errors consistently below 0.3 ms. These errors were constrained by the experimental setup, including the sampling rate of our EEG device and the signal-to-noise ratio affecting pulse detection.

**Significance:** Our timing method achieves sub-millisecond precision in brain-state-dependent closed-loop EEG–TMS, providing a foundation for robust TMS timing that supports adaptive, personalized stimulation protocols. The digital control interface, co-designed with our TMS device, integrates pulse timing and parameters, setting a precedent for future advancements in computer-controlled TMS.

## 1. Introduction

Transcranial magnetic stimulation (TMS) uses pulsed magnetic fields to induce electric fields in the cortex [1,2]. It has shown effectiveness in treating various brain disorders, such as chronic pain and depression [3]. However, current treatment protocols are limited by substantial variability in individual responses and diminishing effectiveness over time [4].

Due to these limitations, interest has grown in individually tailored stimulation protocols, including brain-state-dependent open- and closed-loop protocols [5–8]. For instance, Zrenner et al. [9] raise the possibility that real-time brain-state information could improve treatment outcomes, as motor evoked potentials are strongest when stimulation coincides with the trough (negative peak) of the cortical μ-rhythm, as detected from the subject’s electroencephalogram (EEG).

In brain-state-dependent open-loop TMS, physiological signals, such as EEG, determine the timing and manner of pulse delivery. Similarly, brain-state-dependent closed-loop stimulation uses brain-state information but also incorporates real-time feedback of stimulation effects, thereby “closing the loop” [10]. Both approaches require a fast, reliable pathway between the measuring device and the TMS device to enable accurate and precise stimulation [11].

Signal frequencies relevant for identifying brain states can reach up to 100 Hz [12], with trough-to-peak times as short as 5 ms. Such short-period rhythms demand precise pulse timing, as even a few milliseconds of imprecision can lead to errors in targeting specific signal phases, potentially compromising stimulation effectiveness. Similarly, in cortical μ-rhythm with a center frequency of 10 Hz [13], a 3 ms offset in pulse timing corresponds to a phase error of more than 10°. High precision is also important when investigating unexplored phenomena, where timescales may be unknown; maximizing timing precision helps ensure these effects are accurately captured.

Achieving this level of precision requires TMS devices to support suitable pulse timing methods. Many devices can deliver stimulation pulses with minimal delay in response to Transistor-Transistor Logic (TTL)-level trigger signals [14,15]. While digital messages may introduce delays depending on the communication mechanism [16], they offer greater reliability and compatibility with modern computers [17]. Communication in many other fields has shifted to serial communication-based digital systems over recent decades [18,19], warranting the exploration of digital messaging for timing brain stimulation.

Our work aims to facilitate this transition in TMS. We introduce a method where the TMS device manages pulse timing based on timestamped requests, rather than delivering a pulse immediately upon receiving a request, as is conventional. Building on this method, we present a digital interface for TMS, enabling the use of digital messages to control stimulation timing and other parameters, such as intensity and pulse waveform. Finally, we describe our custom proof-of-concept TMS device that implements this interface, and demonstrate its timing precision.

## 2. Motivation

### 2.1 Robotics and MIDI as Precedents

Before designing an interface for TMS control, it is useful to examine precedents in other fields for insights. In robotics, manufacturers often provide an application programming interface (API) for external control, supporting generic commands such as moving and stopping the robot [20]. Some, including Universal Robots, also offer scripting languages that run directly on the robot hardware [21]. However, pre-determined movement sequences are rarely embedded directly into robots, even for simple, repeatable tasks such as lifting objects, as such hard-coded sequences are typically inadequate for real-world scenarios.

Robotic control offers a compelling analogy for TMS control. Pulse sequences in TMS correspond to movement sequences in robotics, and current simple TMS protocols resemble hard-coded movement sequences. Just as robotics has moved toward flexible, generic control, TMS devices could benefit from adopting similar approaches for complex, adaptive pulse sequences.

Another precedent comes from the external control of electronic musical instruments. In early modular analog synthesizers, sounds were often externally timed using triggers, much like the timing methods of current TMS devices. Later, the introduction of the Musical Instrument Digital Interface (MIDI) standard enabled more sophisticated control of parameters such as timing and pitch [22]. By providing practical methods for synchronization and a shared technological foundation, MIDI significantly advanced computerized music [23].

### 2.2 Limitations of Using Triggers for Timing

Most current TMS devices can deliver stimulation pulses in response to short electrical pulses, typically TTL-level triggers. These binary-valued, continuous-time signals operate at discrete voltage levels, usually 0 and 5 V [24]. The delay between receiving the trigger and delivering the pulse is typically very short, often within a millisecond or less, making triggers a widely used method for precise timing. However, generating these signals from a PC requires a peripheral device, introducing complexity and potential latency.

For example, USB-connected devices in a Linux environment experience approximately 2.5 ms of latency in interrupt transfer mode due to the lack of an optimized USB driver [25]. These latencies, compounded by variable and non-deterministic components, can make achieving precise timing challenging.

Trigger signals also lack sophisticated validation and feedback mechanisms, unlike serial digital communication protocols [26]. This unidirectional nature of triggers prevents the trigger-generating device from confirming whether the TMS device received the signal. Furthermore, the absence of validation means electrical disturbances in the trigger cable could theoretically trigger a pulse, affecting reliability.

Although triggers can manage pulse timing, their limitations become apparent in experimental designs requiring real-time adjustments to stimulation parameters, such as intensity or waveform. For instance, trial-by-trial intensity adjustments require digital messaging, as trigger signals convey only a single bit of information. Combining triggers for pulse timing with serial communication for parameter adjustments increases complexity and the risk of errors. For example, if a digital message to adjust intensity is sent before confirming the delivery of the previous pulse, unexpected delays could cause a race condition—where the timing of concurrent operations leads to unpredictable outcomes [27]—resulting in modification of the incorrect pulse.

### 2.3 Other Pulse Control Methods

Several TMS systems offer USB or RS-232 serial communication interfaces for pulse triggering [28]. While USB can support real-time computing [29], achieving precise timing with these methods presents challenges. The unclear division of timing responsibilities between the control device and its peripherals adds conceptual complexity, increasing the likelihood of programming errors and complicating troubleshooting [30], for instance, by obscuring the cause of delayed pulses.

Modern PCs typically lack built-in serial ports, necessitating USB peripherals or PCI Express extension cards for RS-232 communication. USB peripherals can introduce additional latency due to the USB protocol, while PCI Express extension cards, which offer better timing properties, are typically limited to desktop PCs, as laptop compatibility requires often impractical external adapters. Furthermore, traditional RS-232 communication often has a bandwidth limit of 115,200 bits per second, which constrains the transmission of complex commands to the TMS device.

Latency of these methods remains a significant concern. Triggering via a serial port can introduce delays of up to 10 ms [31], which can be partially mitigated by custom serial port triggering devices connected directly to the TMS device’s trigger input [32]. However, these solutions do not address unpredictable delays originating from the control computer.

Predictability of the control PC can be improved by simplifying its operation. For example, Simulink Real-Time [33] compiles Simulink programs into executables that run outside the operating system (OS) with high predictability (for an example, see Zrenner et al. [9]). However, these setups preclude the use of graphics processing units (GPUs), as GPU functionality requires OS support. This limitation could hinder future stimulation techniques reliant on machine learning algorithms that depend heavily on GPU processing [34–36].

Alternatively, real-time operating systems (RTOSs), such as Real-Time Linux, offer deterministic limits on OS-induced latency, such as task scheduling delays [37]. While RTOSs can improve timing predictability, they do not address delays due to signal transmission or peripheral interfacing. To our knowledge, real-time OS solutions have been rarely applied to TMS, with the exception of NeuroSimo, our open-source platform for closed-loop EEG–TMS [38].

Although sub-millisecond latencies have been demonstrated in simplified setups [32], these results may not extend to real-life scenarios where complex computations on the control PC drive stimulation decisions. Specialized devices can achieve precise trigger timing [39], but similarly to OS-less setups, they generally lack support for GPU-based processing, limiting their applicability for future stimulation algorithms.

### 2.4 Timing Pulses Relative to Feedback Signals

Thus far, we have focused on TMS pulse timing relative to the control PC. Incorporating feedback signals, such as EEG, introduces additional complexity (Figure 1). The total delay *D* from brain measurement to pulse delivery includes deterministic (*D*_*d*_) and stochastic (*D*_*s*_) components [40], where *D* = *D*_*d*_ + *D*_*s*_. For simplicity, we assume *E*[*D*_*s*_] = 0, as any non-zero expected value can be absorbed into *D*_*d*_.

**Figure 1.**
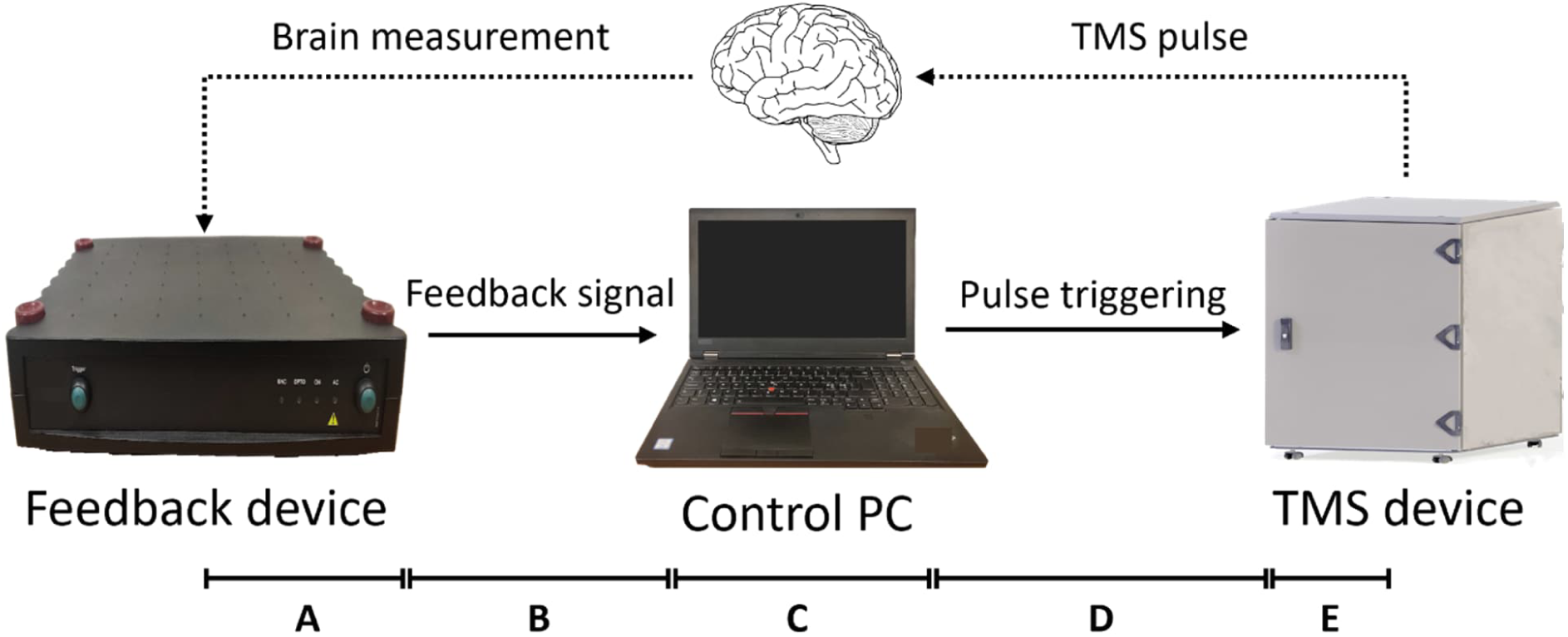
Schematic of a closed-loop stimulation setup. Solid arrows represent direct information flow, while dotted arrows indicate indirect information flow via physical processes such as electromagnetic fields. Assuming negligible delays in brain measurement and pulse delivery, the primary delay components are: (A) Processing delays in the feedback device (e.g., filtering); (B) Transmission delay to the control PC; (C) Processing delays in the control PC (e.g., data preprocessing and timing decisions); (D) Transmission delays due to triggering; (E) Processing delays in the TMS device.

Let us assume a measurement taken at time *t*_0_ results in a decision to deliver a pulse at *t*_0_ + Δ*t*. If the control PC waits for Δ*t*, the pulse is delivered late at *t*_0_ + Δ*t* + *D*, where *D* = *D*_*d*_ + *D*_*s*_ is the total delay. To compensate for the deterministic delay *D*_*d*_, the control PC can instead wait for Δ*t* − *D*_*d*_, delivering the pulse at *t*_0_ + Δ*t* + *D*_*s*_. With *E*[*D*_*s*_] = 0, this approach achieves accurate timing, but precision depends on the variance of *D*_*s*_.

#### Estimating Deterministic Delay

The deterministic delay *D*_*d*_ can be estimated using a fast, direct signaling pathway, such as a trigger signal, from the TMS device to the feedback device. This involves measuring the time difference between a preselected marker in the feedback signal and a return signal from the TMS device, triggered by the marker. The marker is arbitrary, typically corresponding to a specific timepoint in the feedback signal. *D*_*d*_ typically depends on system-specific factors, including the EEG device and its connection to the control PC, the PC’s operating system, and the interaction between the OS and the triggering device. Changes in system properties, such as the OS version or network integrity, can affect *D*_*d*_, introducing a risk of stimulating with outdated delay estimates.

#### Problems with Delay Compensation

Compensating for the deterministic delay introduces several problems:

- Frequent recalibration: Accurate delay estimates may require remeasurement for each stimulation session.
- Gradual degradation: Outdated estimates can gradually degrade stimulation accuracy, which is difficult to detect and troubleshoot [41] and contradicts the fail-fast principle, essential in safety-critical systems such as TMS [42,43].
- Uncompensated random delays: Even with accurate *D_d_* estimates, the stochastic component *D_s_* cannot be corrected, inherently limiting precision.
- Increased complexity: Accounting for delays increases design and operational complexity.

Additionally, achieving strict real-time constraints, where missing deadlines is unacceptable [44], is challenging due to *D*_*s*_, whose variability prevents guaranteeing timely pulse execution.

### 2.5 Real-Time Systems

Real-time systems are designed to process information and complete tasks within defined time constraints. In brain-state-dependent TMS, stimulation pulses are expected to be delivered within specific time windows to ensure efficacy. Each window is centered on the preferred pulse time, with its width determined by the timing precision of the TMS device. While precise timing is important, TMS systems may not demand the strict guarantees of hard real-time systems, where missing a deadline can have catastrophic consequences.

#### Firm vs. Soft Real-Time Constraints

In systems with firm real-time constraints, tasks should meet their deadlines, but occasional missed deadlines do not cause a complete system failure. Soft real-time systems tolerate deviations from deadlines, with utility decreasing gradually rather abruptly. Brain-state-dependent TMS systems likely align more closely with firm real-time requirements, as mistimed pulses may quickly degrade stimulation efficacy, potentially affecting treatment outcomes.

Limited research exists on how mistimed stimulation pulses affect efficacy or safety, particularly in brain-state-dependent open- or closed-loop stimulation. This uncertainty complicates defining precise real-time requirements for TMS systems.

#### Reference Clocks in Real-Time TMS

To deliver pulses within defined time windows, a shared reference clock is essential for establishing a common notion of time. In current EEG–TMS systems, the EEG device clock often serves as the reference, providing timestamps for each sample. However, the TMS device, controlling pulse generation, lacks direct access to the EEG device clock. This disconnect makes the EEG clock a suboptimal choice for the reference—the reference clock should ideally reside in the pulse-generating device.

In a robust real-time TMS system, the TMS device would leverage direct access to the reference clock to ensure precise pulse delivery within defined time windows. To accommodate firm real-time constraints, the device would identify pulses that fall outside the defined window and safely abort them [45]. Additionally, the system should dynamically monitor timing accuracy and relay feedback to the control PC, providing transparency in timing behavior. However, it is difficult to achieve such monitoring and feedback with current TMS systems, which often rely on additional triggers to the EEG device to indirectly provide the feedback. These workarounds add complexity to the system while offering only a limited information channel.

### 2.6 Timing and Synchronization in MIDI

Designing a TMS system to meet firm real-time constraints, such as aborting late pulses, can benefit from examining how similar issues are addressed in MIDI. Both TMS control and MIDI share the requirement of managing external devices with high temporal precision.

MIDI communication is based on messages that describe specific events, such as “note on” or “note off” [46]. In its original form, the MIDI standard only supported immediate execution of transmitted events, comparable to trigger-based TMS control. Later, timestamped messages were introduced, enabling events to be scheduled for playback at precise future times rather than being executed immediately upon receipt [47]. These timestamps are relative to the start of a sequence or song, allowing precise timing that is unaffected by transmission delays or network latency. Furthermore, the close integration of MIDI sound modules with their sound-generating electronics minimizes latency during sound production.

#### Synchronization in MIDI and Its Relevance to TMS

Maintaining synchronization between devices is challenging because their clocks naturally drift out of sync over time due to slight differences in their oscillators [48]. To correct this drift, MIDI relies on periodic clock pulses from a primary device, with each connected device advancing to the next time step only upon receiving a pulse. This mechanism works well for musical timing, where time is divided into relatively large units. However, TMS requires precision in the millisecond range or below, necessitating synchronization methods that support continuous time rather than fixed subdivisions.

In TMS, a distributed clock system can address these requirements. Each device maintains its own clock and uses periodic synchronization pulses to correct drift. Distributed clocks are widely used in protocols such as Network Time Protocol (NTP) and Precision Time Protocol (PTP) [49,50], enabling synchronization with sub-millisecond precision.

#### Problems with Applying Synchronization Protocols to TMS

Although PTP can achieve sub-millisecond synchronization [51], implementing such protocols for TMS presents several problems. First, all system components, including EEG and TMS devices, must support the protocol—a capability typically absent in current EEG devices. Second, achieving high-precision synchronization requires the control PC to use specialized hardware, such as network interface cards with hardware timestamping [52]. These technical barriers highlight the need for synchronization methods tailored specifically for real-time EEG–TMS systems.

### 2.7 Event-Based TMS Control

Inspired by the MIDI standard, an event-based TMS control system can use timed events to transmit simple, discrete commands, such as delivering a single pulse. This approach enables flexible combinations of events, similar to MIDI and robotic control systems, while delegating complex logic to the control PC. By ensuring events are explicit, self-contained, and independent, the TMS device avoids storing or recalling past events, minimizing state management.

In addition to pulse delivery, two other event types—charging and discharging the capacitor— are essential, as they control the stimulation intensity. These events are asymmetric: during a pulse, the capacitor’s energy is discharged and converted into heat, requiring recharging before the next pulse. Over time, capacitor voltage also gradually decreases due to leakage. Additionally, these events operate on different timescales: in our multicoil TMS device [53], a full discharge can take several seconds, whereas charging to maximum voltage typically requires only a few hundred milliseconds. As charging and discharging are physically distinct processes with different durations, treating them as distinct events is logical.

Due to the need to recharge the capacitor between pulses, pulse delivery events could inherently include recharging after the pulse. However, adaptive stimulation introduces complexity that hinders this integration. For example, the intensity of the next pulse may not yet be determined when the previous pulse is delivered, and long intervals between pulses can lead to additional recharging due to voltage leakage. Therefore, it is more efficient to keep event types simple and distinct, avoiding compound events such as pulse delivery followed by recharging.

Proper handling of concurrent events is essential in event-based designs. For example, delivering a pulse while simultaneously charging the capacitor is undesirable, as the voltage change during charging can interfere with pulse generation. An event-based TMS control system should, therefore, prohibit concurrent events that affect capacitor voltage, ensuring the integrity of stimulation.

An additional event type is inspired by the capability of many TMS devices to transmit TTL-level triggers concurrently with stimulation pulses, signaling pulse timing to external devices. Similarly, gate signals are often generated to indicate ongoing pulses (Figure 2). Both triggers and gates can be implemented through a generalized “binary signal out” event, which specifies the start time and duration: the TMS device sets the TTL-level signal to high voltage for the specified duration, then returns it to low voltage. For example, a trigger might be represented as a brief pulse (e.g., 20 μs), while the gate duration corresponds to the pulse length. Gates can also be programmed to start before and extend beyond the pulse. These flexible triggers and gates, which can be timed independently of pulse events, are particularly useful in research applications where custom devices are integrated with the TMS system.

**Figure 2.**
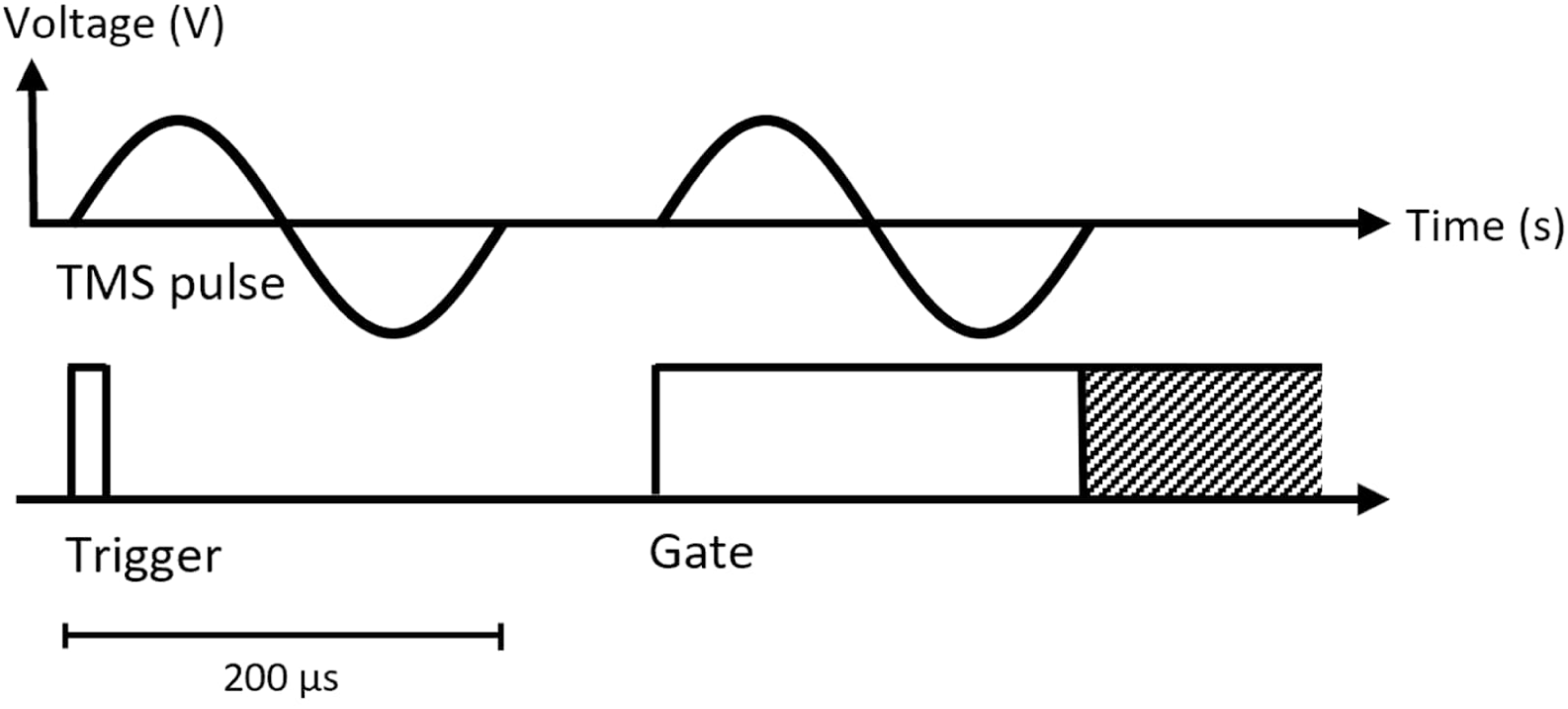
TMS pulses (top) with simultaneous trigger and gate signals (bottom). The trigger signal (left) marks the pulse start, while the gate signal (right) lasts for the pulse duration or longer (shaded area). Trigger signals communicate pulse timing to external devices, such as EEG systems, while gate signals can prevent data collection during pulses, avoiding TMS-induced artifacts.

## 3. Design

This section describes a comprehensive timing and control system for TMS devices, designed to enable high-precision stimulation in closed-loop EEG–TMS setups.

We begin by presenting a TMS timing method that utilizes timestamped messages, addressing the limitations of immediate pulse delivery methods, such as timing variability and problems with meeting strict real-time constraints.

Next, we extend this method to brain-state-dependent, closed-loop EEG–TMS using a distributed clock system inspired by MIDI, eliminating the need for explicit latency compensation.

Finally, we propose a versatile TMS control interface that supports timestamped messaging, detail its implementation in a proof-of-concept TMS device, and demonstrate its precision in aligning pulses with EEG signals.

### 3.1 Timestamped Messages for TMS Control

Our proposed timing method relies on serial digital messages specifying precise execution times for pulses: the TMS device maintains an internal clock, waits until the designated time, and then generates the pulse. This contrasts with traditional methods, where a pulse is delivered immediately upon receiving either a TTL-level trigger or a digital message (Figure 3).

**Figure 3.**
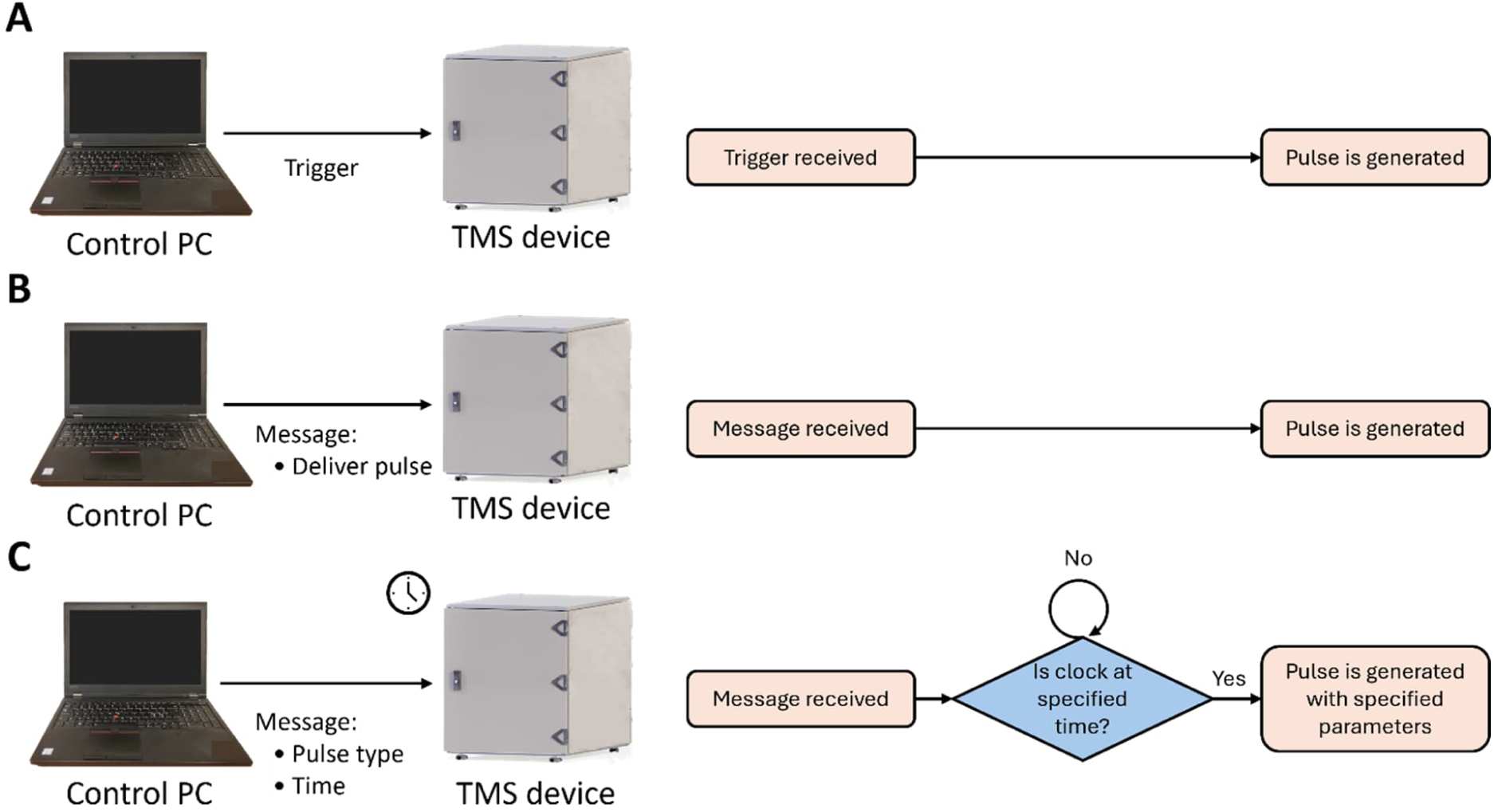
Comparison of pulse timing methods: (A) TTL-level trigger, (B) digital message for immediate pulse delivery upon receipt, and (C) digital message with timing information. In (C), the TMS device uses an internal clock (indicated by the clock symbol) to wait until the specified time before generating the pulse, ensuring precise timing. The message in (C) can also specify pulse parameters, such as intensity and type.

Timestamped commands enable externally defined pulse sequences, managed by the TMS device based on its internal clock. This allows for efficient hardware-based pulse timing, achieving theoretical precision down to a single clock cycle. Additionally, the TMS device can abort a pulse if it cannot be executed within its designated time window, supporting firm real-time constraints.

By combining pulse timing and parameters into a single message, timestamped commands eliminate the need for separate communication channels, which traditional methods would suggest, using triggers for precise timing and digital messaging for parameter setting. This single-channel approach reduces communication complexity and avoids potential coordination issues.

Like conventional methods, timestamped messaging supports adaptive stimulation that responds to feedback signals, such as EEG. For instance, stimulation intensity can be dynamically adjusted based on real-time EEG signal power. However, brain-state-dependent stimulation, such as targeting specific EEG signal phases, requires latency compensation, as the EEG and TMS devices operate on separate clocks.

### 3.2 Extension for Brain-State-Dependent EEG–TMS

Our timing method extends to brain-state-dependent, closed-loop EEG–TMS systems, where TMS pulses must be precisely timed relative to the EEG signal. This extension is enabled through a distributed clock system (see Section 2.6), with the TMS device designated as the primary device, as discussed in Section 2.5. The primary device periodically transmits synchronization pulses to maintain clock alignment across devices (Figure 4).

**Figure 4.**
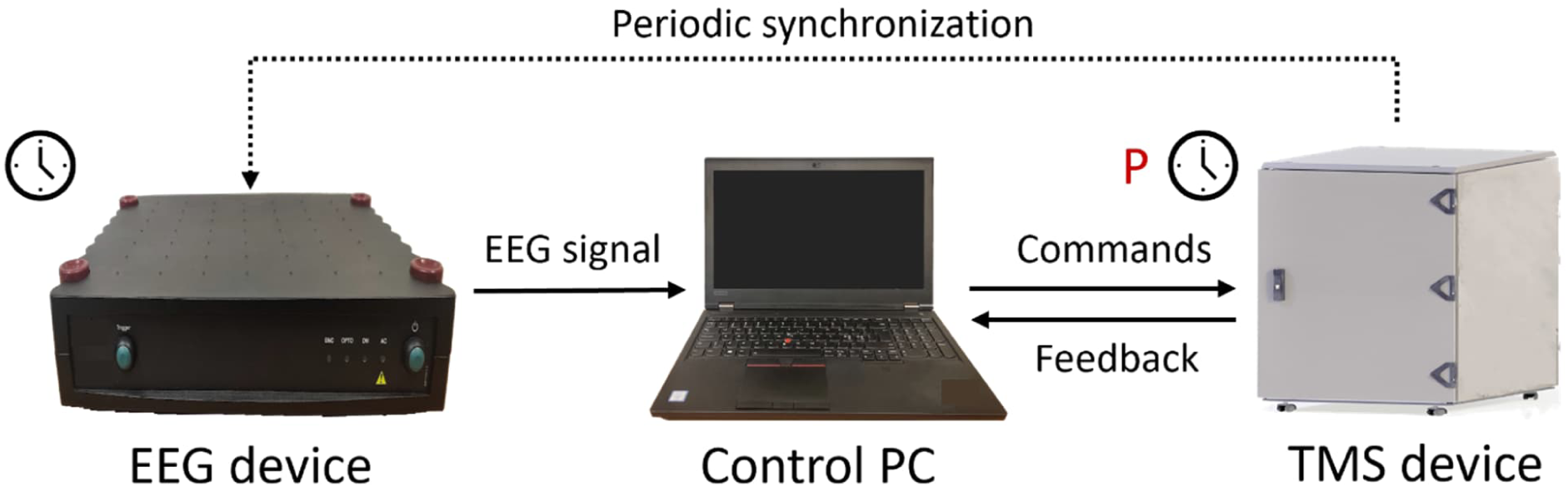
Device setup and synchronization. Arrows represent information flow. Independent clocks (indicated by clock symbols) are maintained by the EEG and TMS devices, with the TMS device serving as the primary clock (P). The TMS device periodically transmits synchronization pulses to the EEG device, allowing the control PC to correct EEG timestamps using these pulses.

To address the lack of native support for clock synchronization protocols in EEG devices, our method uses two key insights:

1. Incidental synchronization pulses: EEG devices typically support receiving triggers, which can be used as synchronization pulses, although they are not inherently designed for that purpose.
2. Independent operation of the control PC: Precise pulse timing does not require the control PC to synchronize directly with the EEG and TMS devices.

In our approach, the control PC corrects clock drift by analyzing the synchronization pulses, which are timestamped by the EEG device. This enables the control PC to calculate deviations from the expected intervals between consecutive pulses and adjust the timestamps of subsequent EEG samples to ensure alignment.

The theoretical clock alignment achievable with this method is constrained by the maximum drift between synchronization pulses. In our reference implementation, synchronization pulses are transmitted at 1-second intervals, achieving microsecond-level precision assuming a typical drift rate of 1 part per million [54]. High timing precision is achievable even with lower EEG sampling rates, as many EEG devices can timestamp triggers with microsecond-precision regardless of the sampling rate (e.g., see Bittium NeurOne manual [55]).

### 3.3 Digital Interface for TMS Control

Building on the timestamped messaging approach, we designed an event-based digital interface for TMS control. This interface supports both traditional pre-defined pulse sequences without EEG feedback, as well as brain-state-dependent open- and closed-loop EEG–TMS.

The interface defines several event types, including pulse delivery, stimulation intensity adjustment, and TTL-level output signal generation. Events can be scheduled for immediate execution, at specified future times, or triggered externally, as in conventional methods— allowing combinations such as delivering pulses with simultaneous TTL-level output signals.

To establish a consistent notion of time, TMS operation is divided into sessions, with the session start serving as the zero-time reference for all subsequent timing. At the end of a session, the TMS device halts its internal clock, prevents further event requests, and fully discharges the capacitor to ensure a well-defined initial state for the next session.

Safety mechanisms prohibit concurrent events that could compromise voltage integrity, such as simultaneous pulse delivery and capacitor charging or discharging. However, other concurrent actions, such as delivering stimulation pulses alongside TTL-level output signals, are permitted, providing flexibility.

#### Event Messages

Event request messages include the following fields:

- Event ID: A unique unsigned integer identifying the event.
- Event type: Specifies the type of event—pulse, charge, discharge, or signal.
- Execution condition: Defines when the event occurs—immediate, timed, or triggered externally.
- Execution time (optional): A floating-point number specifying the execution time for timed events.

Event IDs uniquely identify events within a session and are referenced in feedback messages sent to the control PC upon event completion. Each feedback message includes the event ID, start and finish times, and an error code—0 if the event was successful. Additional fields specific to the event type are also included, such as the target voltage for charging or discharging, or the duration for output signal events.

#### Control Commands and Queries

The interface supports the following control commands:

- Start session: Begins a new session, resetting the device state and internal clock.
- End session: Ends the current session, fully discharges the device, and stops the internal clock.
- Abort event: Aborts a future or ongoing event, identified by its event ID.

State variables of the TMS device can also be queried, including:

- Current time: The current value of the device’s internal clock.
- Capacitor voltage: The current voltage of the capacitor.
- Session status: The current session state (active or inactive).
- Error status: The operational status code (0 for normal operation).

Additional queries include the time required to charge or discharge the device to a specified voltage, the maximum supported voltage, the device model, and the firmware version.

Figure 5 presents an example session for a simple repetitive TMS (rTMS) experiment.

**Figure 5.**
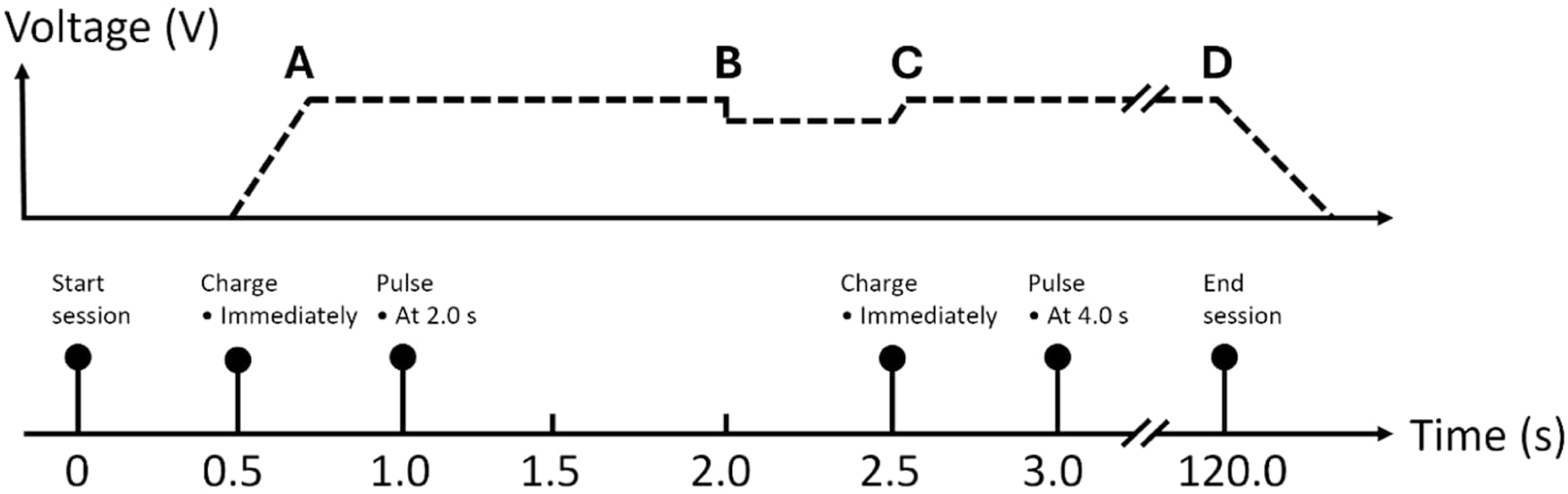
Example timeline of a simple rTMS experiment with two-second intertrial intervals. The capacitor voltage (top, dashed line) changes over time due to periodic charge and pulse events (bottom). (A) The capacitor is charged immediately in response to a charge event. (B) A pulse is delivered at 2.0 seconds. (C) The capacitor is recharged. (D) The session concludes, and the capacitor is fully discharged.

### 3.4 Extension for Multicoil TMS

Multicoil TMS enables more flexible stimulation patterns by simultaneously using multiple stimulation coils [56,57]. A multicoil TMS device can be realized by equipping the device with multiple independent pulse generators, each controlling a single coil, as done in our multicoil TMS device [53].

To support multicoil functionality, the interface extends event request messages with a target coil field, specifying the coil for which the event is intended. Each coil control unit processes only the events directed to it, ensuring modularity and scalability. Concurrency rules are adapted to the multicoil setup:

- Simultaneous pulses across multiple coils are permitted.
- Concurrent discharging of multiple control units is allowed.
- Concurrent charging is supported if each unit has its own dedicated charger.

### 3.5 Proof-of-Concept Implementation

To evaluate the interface, we developed a multicoil TMS device and conducted technical characterization as well as cortical mapping experiments using the interface [58]. Additionally, we created MATLAB and Python APIs to facilitate control software development and scripting for the device (Appendix A).

The device is built around a field-programmable gate array (FPGA) core provided by National Instruments (NI). The FPGA and control PC communicate via Thunderbolt, exchanging serialized messages through an NI-provided C++ module. The FPGA manages the internal clock, processes incoming event requests, and controls the device electronics accordingly. When an EEG device is connected, the FPGA also transmits periodic synchronization pulses via a TTL-level output port.

#### Application Examples

To demonstrate the versatility of the proposed interface, we implemented common stimulation patterns as practical examples. Pseudo-code for these examples is included in Appendix B.

1. *Repetitive TMS with Simultaneous Trigger Out*

In repetitive TMS (rTMS), periodic pulse generation alternates with capacitor charging. Simultaneously with each pulse, a short signal-out event produces a trigger signal, indicating pulse timing. The session begins before the sequence and ends after its completion, ensuring proper initialization and discharging of the capacitor.

2. *Chaining TMS Devices for Paired-Pulse Stimulation*

Two TMS devices, designated as leader and follower, can be chained by connecting the signal-out port of the leader to the trigger-in port of the follower. This setup enables paired-pulse stimulation, where the leader generates the first pulse and a delayed trigger signal, prompting the follower to generate the second pulse. This mirrors paired-pulse functionality found in commercial systems such as the Magstim BiStim^2 1^.

3. *EEG-Guided TMS*

In EEG-guided stimulation, EEG data are buffered and processed by a classifier that determines whether and when to stimulate, based on drift-corrected timestamps. With efficient implementation, 99% of stimulation pulses can be timed within 6.5 milliseconds of acquiring the latest EEG sample when targeting the trough of μ-rhythm [38].

### 3.6 Experimental Verification

To validate the precision of our pulse timing method, we conducted an EEG–TMS experiment where TMS pulses were delivered periodically, timed relative to the EEG signal. Timing errors were quantified by comparing theoretical pulse times to TMS-induced pulse artifacts recorded in the EEG, which served as proxies for the actual pulse timing.

Over a 10-minute session, 600 single pulses were delivered at 0.99-second intervals. This slightly offset interval avoided alignment with synchronization pulses, transmitted once per second, simulating a realistic non-synchronized scenario. A bipolar EEG electrode placed at the center of the stimulation coil registered pulse artifacts, while a reference electrode was positioned farther away. Simultaneous trigger signals transmitted to the EEG device provided a direct measure of timing precision between TMS and EEG (Figure 6).

**Figure 6.**
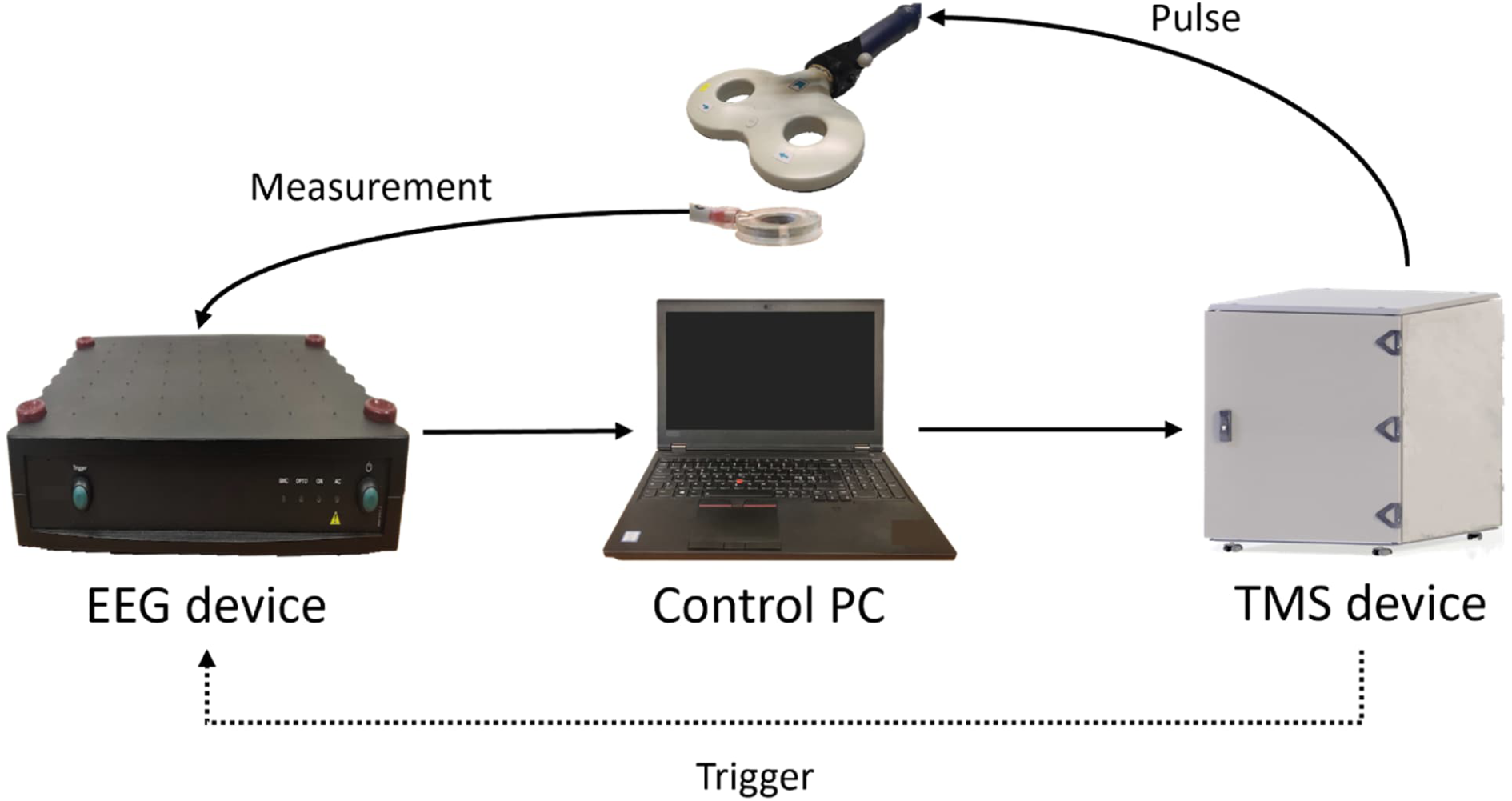
Experimental setup for verifying pulse timing precision. TMS pulses were delivered at 0.99-second intervals to avoid alignment with synchronization pulses. A bipolar electrode at the center of the TMS coil registered pulse artifacts, while simultaneous trigger signals transmitted to the EEG device (dotted line) provided a direct timing measure.

EEG data were recorded using Bittium NeurOne (Bittium Plc, Finland) at a sampling rate of 5 kHz, yielding a 0.2 ms sampling period. Data stream was sent to the control PC, where raw timestamps were corrected for drift (see Section 3.2). Both the drift-corrected timestamps and voltage measurements from the electrode were stored for subsequent analysis.

#### Pulse Detection and Timing Analysis

Pulse artifacts were detected from the EEG signal by differencing the voltage time series, which served as a simple edge-detector, and applying a threshold to the differenced series. The latter sample of each difference exceeding the threshold was designated as the pulse time. Binary search was used to determine a threshold that detected exactly 600 pulses. Pulses detected less than 0.1 seconds apart were excluded to avoid multiple detections of the same pulse.

Our pre-hoc theoretical analysis identified two factors that constrain the maximum achievable timing precision with this method: the sampling period *T* of the EEG device and the minimum pulse detection delay *T*_*d*_, caused by the limited signal-to-noise ratio of the measurements (Figure 7). Together, these factors limit the maximum precision to *T*+ *T*_*d*_.

**Figure 7.**
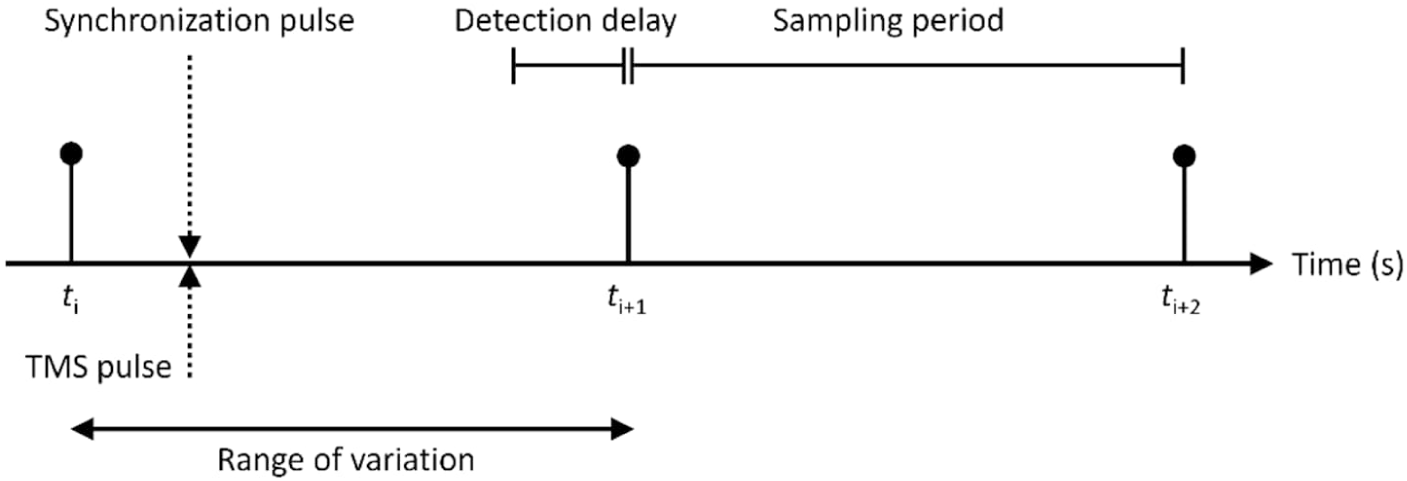
Temporal relationship between a synchronization pulse (dotted downward arrow), simultaneous TMS pulse (dotted upward arrow), and periodic EEG sampling (pins at *t*_*i*_, *t*_*i*+1_, and *t*_*i*+2_). Clock drift shifts the relative timing of the synchronization pulse, and thus the TMS pulse, across its range of variation (solid line with arrows at both ends). When a stimulation pulse occurs between *t*_*i*_ and *t*_*i*+1_, it is typically detected at *t*_*i*+1_. However, if the pulse occurs after *t*_*i*+1_ − *T*_*d*_ (where *T*_*d*_ is the minimum pulse detection delay caused by limited signal-to-noise ratio), noise prevents immediate detection, delaying it to *t*_*i*+2_. In the worst-case scenario, the pulse occurs precisely at *t*_*i*+1_ − *T*_*d*_, resulting in detection at *t*_*i*+2_. Thus, the maximum delay between the pulse and its detection is *T*+ *T*_*d*_, where *T* is the sampling period.

Triggers, however, provide a more precise estimate of timing alignment between the TMS and EEG devices. Unlike pulses, triggers are detected immediately, and their timestamps with NeurOne are not limited by the EEG sampling rate. However, the precision of trigger timing does not fully reflect the precision of pulse delivery, as trigger and pulse generation follow separate electronic pathways in the TMS device.

To investigate this discrepancy, we measured the timing difference between triggers and actual pulses by connecting both the trigger output and a probe coil detecting pulses to an oscilloscope. This setup enables us to assess the trigger-to-pulse error and evaluate the reliability of using trigger timing as a proxy for pulse precision.

## Results

All 600 pulses were successfully detected using the threshold criterion, with no false positives. Timing errors relative to the theoretical pulse times were less than 0.3 ms for every pulse, consistent with the limits set by the sampling period (0.2 ms) and an unknown pulse detection delay. Using Laplace’s rule of succession, the Bayesian probability that the next pulse will have a timing error smaller than 0.3 ms exceeds 99.8%, demonstrating high timing precision.

Over the 10-minute experiment, the uncorrected EEG and TMS clocks drifted apart by 3.9 ms (averaging 6 µs per second). This confirms the necessity of drift correction to maintain accurate timing, especially for experiments exceeding 30 minutes. The observed drift was linear (not shown), supporting the use of regular drift corrections to keep errors within acceptable bounds.

The trigger timing error remained below 6.0 µs for all 600 triggers, constrained by the drift between synchronization pulses. The mean trigger-to-pulse timing error was 2.9 µs, with a minimal standard deviation of 0.04 µs, demonstrating consistent alignment between pulses and simultaneous triggers.

These results confirm that the proposed method achieves precise TMS pulse timing relative to EEG signals, with errors below 0.3 ms. Additionally, the microsecond-scale trigger timing and trigger-to-pulse alignment suggest that pulse timing precision can surpass the limits imposed by the present experimental design, potentially achieving microsecond-level precision.

## 4 Discussion

We introduced a timing method and digital interface for TMS control, achieving sub-millisecond precision in closed-loop EEG–TMS. This method transitions from reactive, synchronous TTL-level triggers to proactive, asynchronous pulse sequence planning, aligning with the evolving demands of adaptive stimulation.

By delegating timing responsibility to TMS device hardware, our method ensures that pulses are either delivered on time or safely discarded, supporting firm real-time properties. Timing precision can potentially reach clock cycle resolution, constrained only by the electronic pathways for pulse generation. Managing timing directly within the device electronics eliminates vague divisions of timing responsibilities between system components.

Extending this method to closed-loop EEG–TMS through synchronized TMS and EEG device clocks enables precise, feedback-driven stimulation without requiring delay compensation in the signal pathway. Importantly, variability in signal pathway delays does not affect timing precision, simplifying system design and providing a robust framework for closed-loop EEG–TMS.

Our digital interface was co-designed with the TMS device to optimally support external control, reflecting the broader shift from hardware to software functionality [59–61] and the principles of hardware-software co-design [62,63]. It includes features valuable for EEG–TMS [64], such as adjustable recharge delays and support for different pulse waveforms. Its minimalist design facilitates the creation of complex stimulation sequences from simple, well-defined operations. This design philosophy reduces hardware complexity and enables advanced logic to be managed by the control PC.

While our interface could be adapted to other TMS devices with minor modifications, certain design decisions are tailored to our multicoil TMS device. Additional features, such as querying pulse duration, may improve general applicability, especially as most TMS devices use internally predefined waveforms rather than allowing arbitrary ones.

Currently, the interface requires engineering expertise, making it most suitable for research applications. Future iterations could include a software layer with a graphical user interface (GUI) for clinicians, streamlining tasks such as pulse sequence planning or selecting adaptive stimulation algorithms.

The interface offers significant potential for future extensions. For instance, soft real-time constraints could be supported by optionally allowing late pulse delivery. Additionally, compound events, such as paired-pulse generation, could simplify defining complex stimulation sequences, reducing the need for multiple individual events.

Pausing sessions is not directly supported, as this would require halting the clocks of both the TMS and EEG devices, complicating synchronization. Instead, sessions can effectively be paused by halting event requests from the control PC, offering a practical workaround.

Our design separates charging and discharging into distinct events, requiring the control PC to monitor capacitor voltage and determine the appropriate action, adding operational complexity. Stimulation strength is currently defined in terms of capacitor voltage, whereas many commercial TMS devices use the percentage of maximum stimulator output (MSO). However, both MSO and capacitor voltage are difficult to interpret directly in terms of brain effects. Future work should explore more robust measures of stimulation strength, such as electric field strength, which may offer a clearer link to biological effects.

Future research should also explore the adoption of existing protocols, such as Precision Time Protocol (PTP), for synchronizing devices in closed-loop EEG–TMS. The lack of PTP support in current TMS and EEG devices increases synchronization complexity. Ideally, other devices, such as robotic systems for moving the stimulation coil [65], would also share the synchronized clock. This would enable high-precision coordination of robot movements with stimulation pulses, allowing dynamic protocols where stimulation coil physically moves during a session.

Advances in pulse-width modulation (PWM) of stimulation waveforms [66,67] could simplify stimulator output modification by eliminating the need for charging or discharging to adjust intensity between pulses, as PWM can account for varying intensity. However, PWM introduces challenges, such as managing complex pulse waveforms. Future work should explore effectively integrating these advances with the current control interface.

The extent of real-time constraints necessary for adaptive TMS is not yet fully defined. However, adopting the theoretical framework of real-time systems provides a grounded basis for establishing these requirements, regardless of their level, and defining acceptable timing tolerances for various stimulation protocols. This approach can lead to clearer standards that account for both safety and clinical efficacy.

In this article, we presented a timing method and control interface for TMS, offering a robust solution for precise pulse timing in brain-state-dependent, closed-loop EEG–TMS. As TMS functionality continues to shift from hardware to software, we anticipate significant advancements in the field, enabling the development of versatile and robust TMS interfaces similar to ours. These advancements will support the transition from simple pulse sequences to adaptive, complex protocols, ultimately improving treatment outcomes.

## Acknowledgments

This work has been supported by the European Research Council (ERC Synergy) under the European Union’s Horizon 2020 research and innovation programme (ConnectToBrain; grant agreement No 810377) and the Finnish Cultural Foundation. We thank Tuomas J. Lukka, Tomi-Mikael Kahilakoski, and Veli Peltola for useful discussions, Victor H. Souza for comments on the manuscript, and Robin Rantala for the model picture of the multicoil TMS device. The image of the human brain in lateral view (source: Wikimedia Commons, originally from https://www.rawpixel.com/image/6289600/) was used in accordance with its public domain dedication (CC0 1.0 Universal). OpenAI’s GPT-4 and Anthropic’s Claude 3.5 Sonnet were used as tools for revising the manuscript. All final content has been reviewed and approved by the authors.

## Author Contributions

Olli-Pekka Kahilakoski: Conceptualization, Methodology, Software, Validation, Formal Analysis, Writing – original draft, reviewing & editing. Heikki Sinisalo: Conceptualization, Methodology, Validation, Writing – reviewing & editing. Jaakko O. Nieminen: Conceptualization, Methodology, Writing – reviewing & editing. Kyösti Alkio: Software, Writing – reviewing & editing. Kim Valén: Software, Writing – reviewing & editing. Gábor Kozák: Conceptualization, Writing – reviewing & editing. Risto J. Ilmoniemi: Project administration, Supervision, Funding acquisition, Writing – reviewing & editing. Timo Roine: Conceptualization, Project administration, Supervision, Writing – reviewing & editing.

## Data Availability Statement

The dataset and code supporting this study are openly available in Zenodo at https://doi.org/10.5281/zenodo.14198232.

## Supplementary Materials

## Appendix A Python API Description

This appendix describes the Python API functions available for device control and experiment management.

### Device Control Functions

**`start_device()`**

Start the device.

**`stop_device()`**

Stop the device.

`get_device_state()`

Return the current state of the device.

**Return value**

- `DeviceState.NOT_OPERATIONAL` – Device is unoperational
- `DeviceState.STARTUP` – Device is starting up
- `DeviceState.OPERATIONAL` – Device is operational
- `DeviceState.SHUTDOWN` – Device is shutting down

### Experiment Control Functions

`start_experiment()`

**Start the experiment**.

`stop_experiment()`

**Stop the experiment**.

`get_experiment_state()`

Return the current state of the experiment.

**Return value**:

- ExperimentState.STOPPED – Experiment is stopped
- ExperimentState.STARTING – Experiment is starting
- ExperimentState.STARTED – Experiment is started
- `ExperimentState.STOPPING` – Experiment is stopping

### Event and Measurement Functions

`get_event_feedback(id)`

Return the feedback for the specified event.

**Parameters**:

- `id` (int) – Event identifier (range: 0–65535)

**Return value**:

- Integer feedback value if available, `None` otherwise

`get_voltage(channel)`

Return the capacitor voltage.

**Parameters**:

- `channel` (int) – Channel number (range: 1–5)

**Return value**:

- Voltage in volts (V)

`get_pulse_count(channel)`

Return the total number of pulses generated.

**Parameters**:

- `channel` (int) – Channel number (range: 1–5)

**Return value**:

- Number of pulses

`get_temperature(channel)`

Return the coil temperature.

**Parameters**:

- `channel` (int) – Channel number (range: 1–5)

**Return value:**

- Temperature if sensor is present, `None` otherwise

`get_time()`

Return the current time from the start of the session.

### Event Request Functions

`request_charge(event_id, execution_condition, time, channel, target_voltage)`

Request a charging of the capacitor.

**Parameters**:

- `event_id` (int) – Event identifier (range: 0–65535)
- `execution_condition` (ExecutionCondition) – One of:

- ExecutionCondition.IMMEDIATE – Execute immediately
- `ExecutionCondition.TIMED` – Execute at specified time
- ExecutionCondition.TRIGGERED_EXTERNALLY – Execute on external trigger
- `time` (float) – Execution time (used only with `ExecutionCondition.TIMED`)
- `channel` (int) – Channel number (range: 1–5)
- `target_voltage` (float) – Target voltage (range: 0–1500 V)

`request_discharge(event_id, execution_condition, time, channel, target_voltage)`

Request a discharging of the capacitor.

**Parameters**:

- `event_id` (int) – Event identifier (range: 0–65535)
- `execution_condition` (ExecutionCondition) – One of:

- ExecutionCondition.IMMEDIATE – Execute immediately
- `ExecutionCondition.TIMED` – Execute at specified time
- ExecutionCondition.TRIGGERED_EXTERNALLY – Execute on external trigger
- `time` (float) – Execution time (used only with `ExecutionCondition.TIMED`)
- `channel` (int) – Channel number (range: 1–5)
- `target_voltage` (float) – Target voltage (range: 0–1500 V)

`request_pulse(event_id, execution_condition, time, channel, waveform)`

Request a pulse.

**Parameters**:

- `event_id` (int) – Event identifier (range: 0–65535)
- `execution_condition` (ExecutionCondition) – One of:

- ExecutionCondition.IMMEDIATE – Execute immediately
- `ExecutionCondition.TIMED` – Execute at specified time
- ExecutionCondition.TRIGGERED_EXTERNALLY – Execute on external trigger
- `time` (float) – Execution time (used only with `ExecutionCondition.TIMED`)
- `channel` (int) – Channel number (range: 1–5)
- `waveform` (list) – List of dictionaries with keys:

- ’`mode`’: One of:

- PulseMode.RISING
- PulseMode.HOLD
- PulseMode.FALLING
- PulseMode.ALTERNATIVE_HOLD
- ’`duration_in_ticks`’ (int): Range 0–65535

`request_signal_out(event_id, execution_condition, time, port, duration_us)`

Request a TTL-type output signal.

**Parameters**:

- `event_id` (int) – Event identifier (range: 0–65535)
- `execution_condition` (ExecutionCondition) – One of:

- ExecutionCondition.IMMEDIATE – Execute immediately
- `ExecutionCondition.TIMED` – Execute at specified time
- ExecutionCondition.TRIGGERED_EXTERNALLY – Execute on external trigger
- `time` (float) – Execution time (used only with `ExecutionCondition.TIMED`)
- `port` (int) – Port number (range: 1–2)
- `duration_us` (int) – Signal duration in microseconds (range: 0–65535)

`wait_for_completion(timeout=None)`

Wait for the completion of all requested events.

**Parameters**:

- `timeout` (float, optional) – Maximum wait time. If not specified, waits indefinitely.

## Appendix B: Example Implementations in Pseudo-code

This appendix provides pseudo-code implementations of the stimulation protocols described in the main text.

### Repetitive TMS with Synchronous Trigger Out

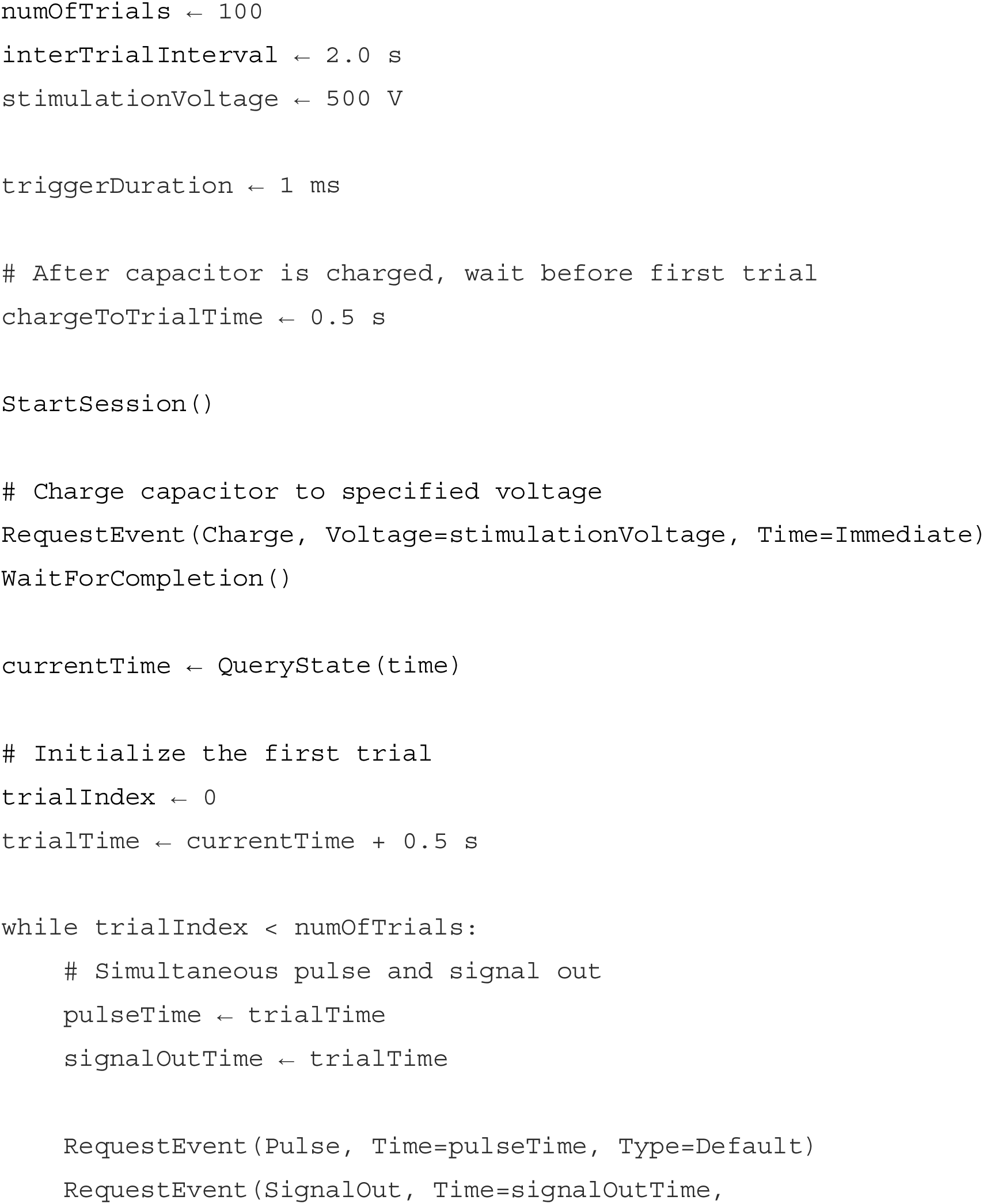
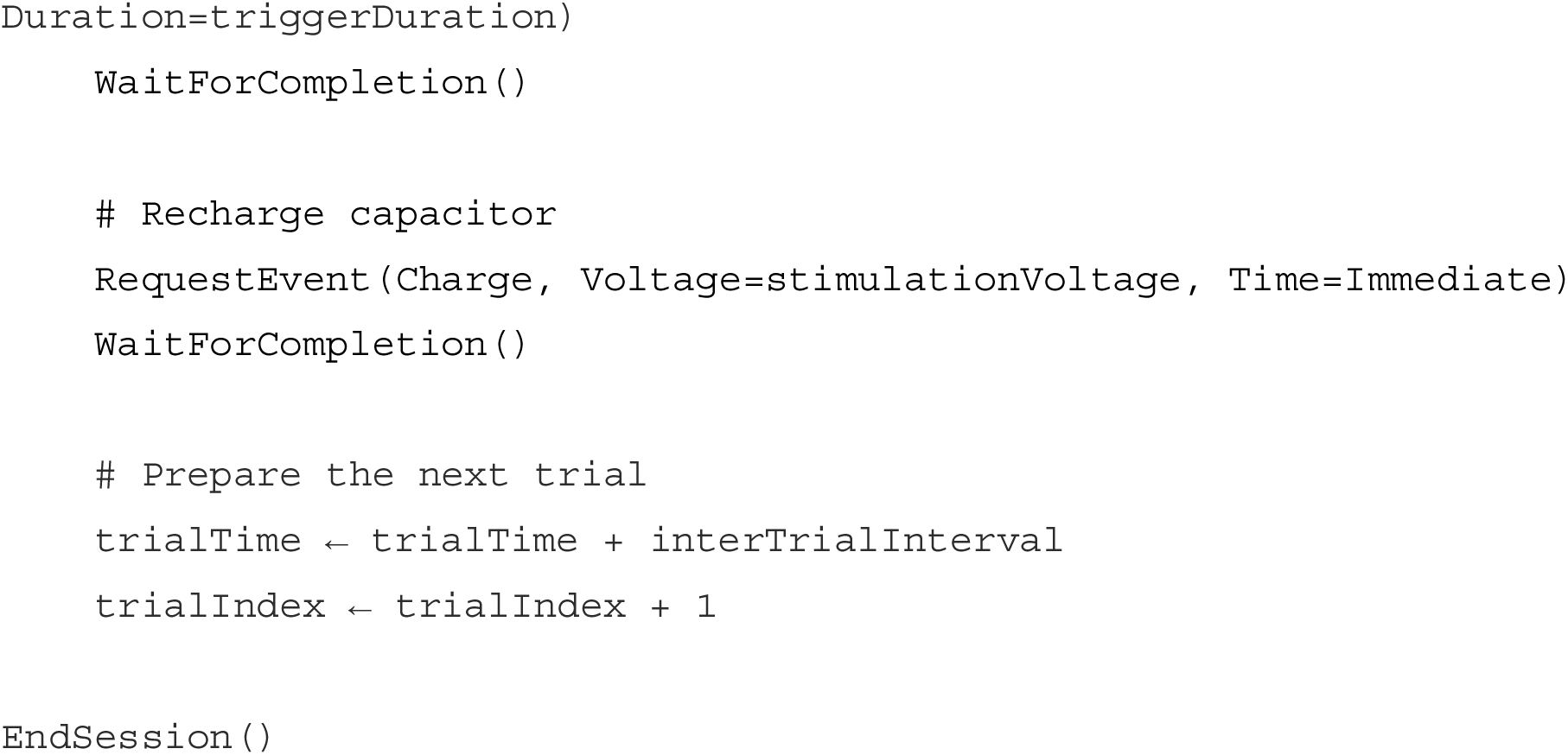

### Chained Paired-Pulse TMS

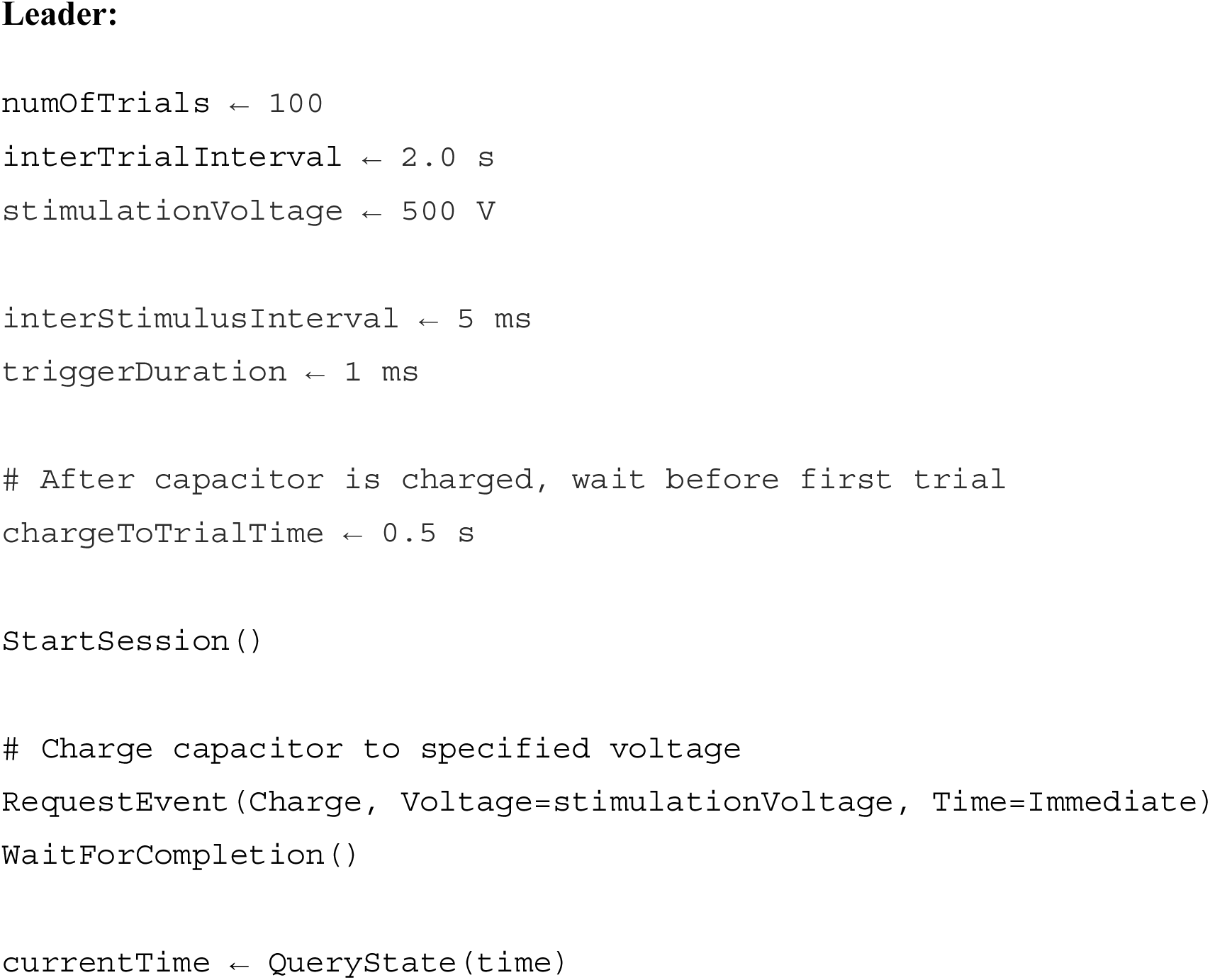
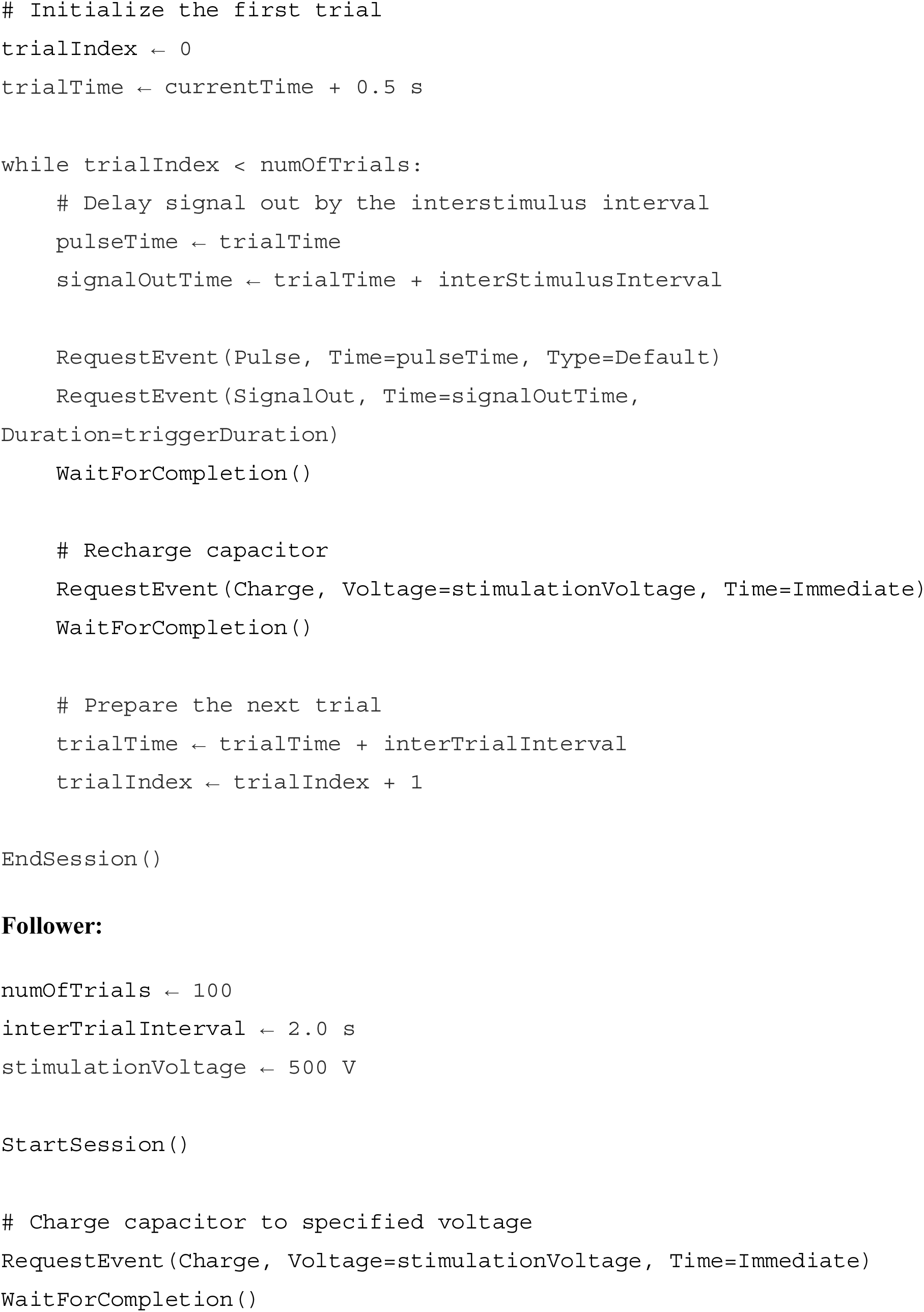
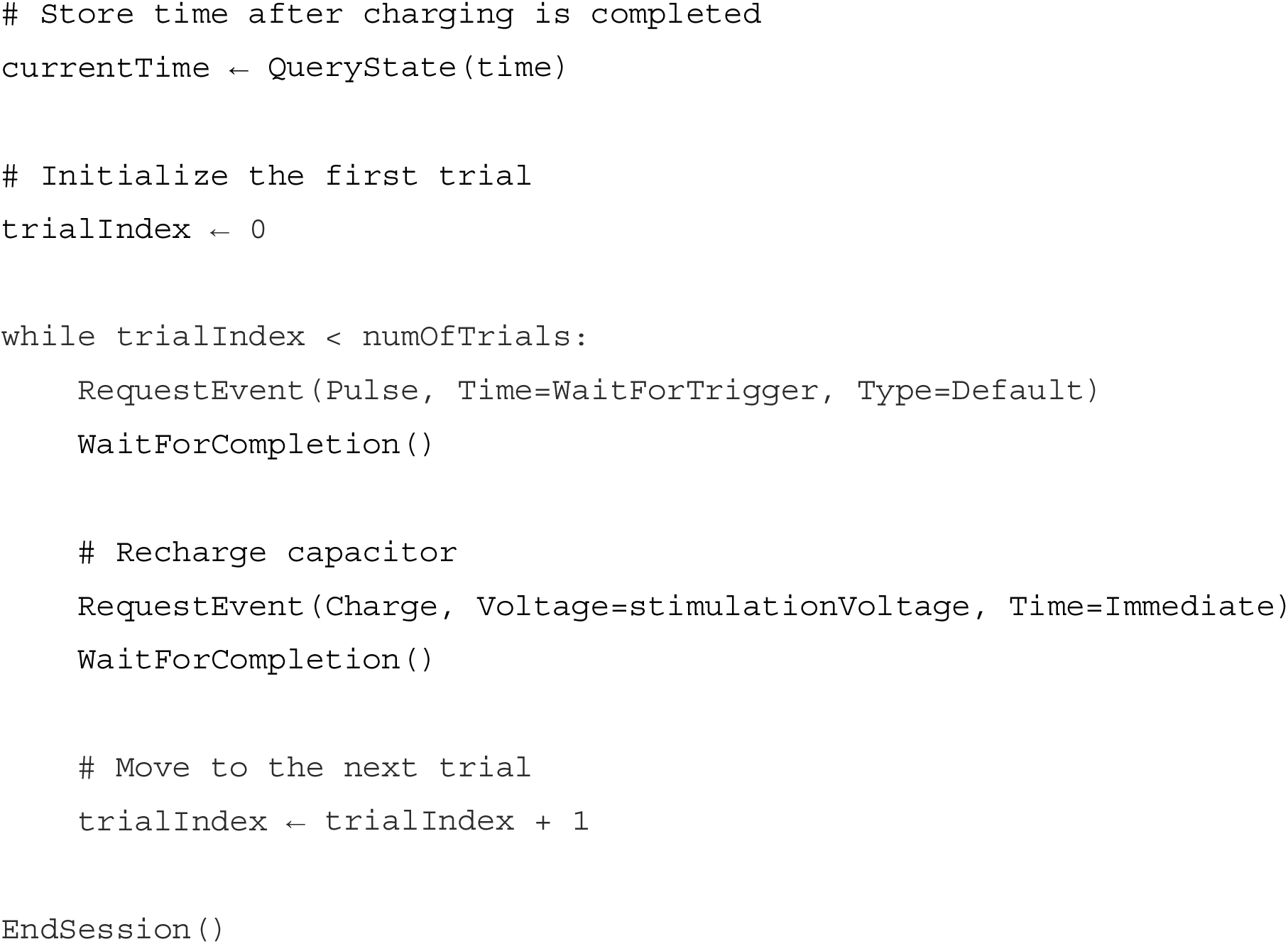

### EEG-Guided Stimulation

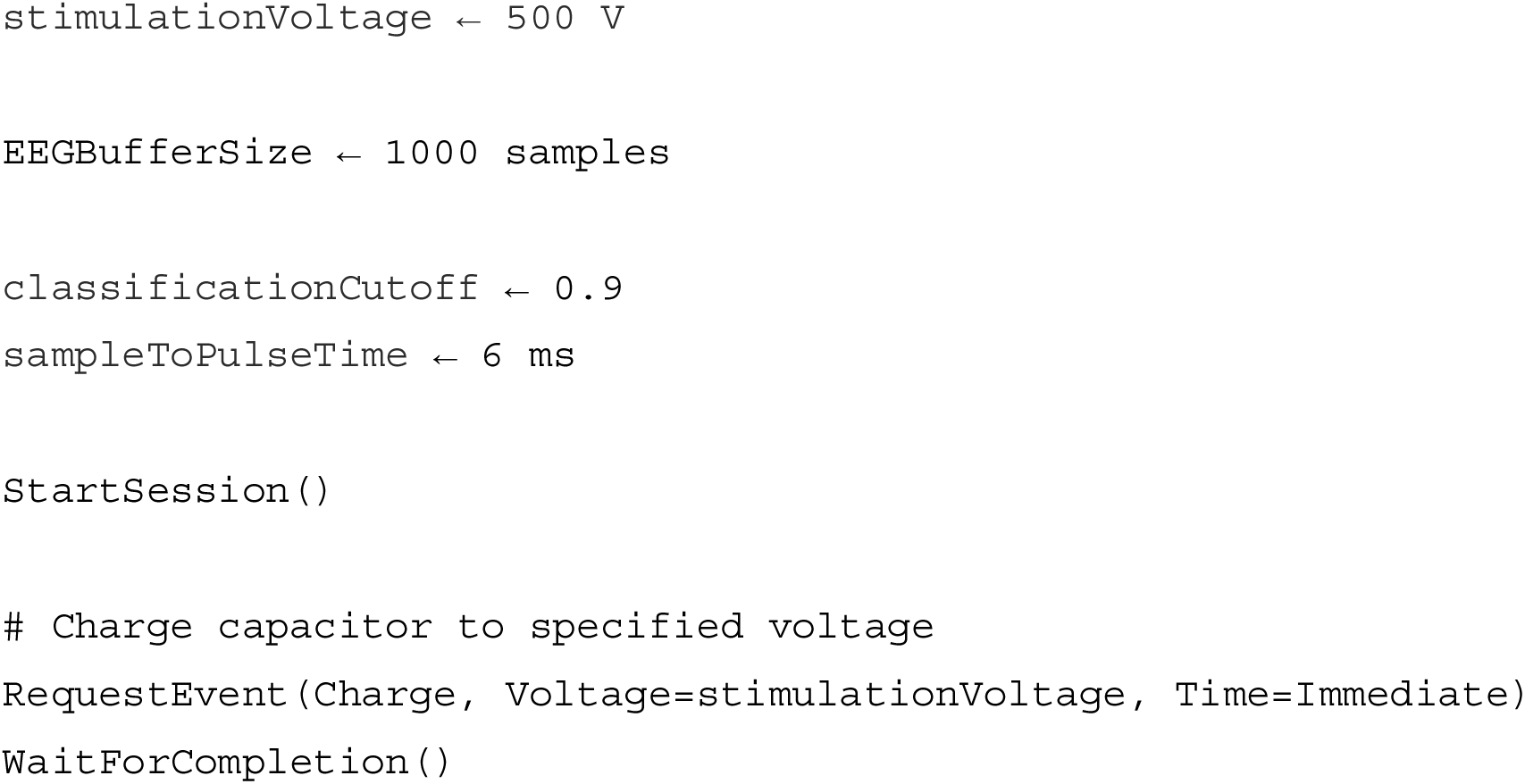
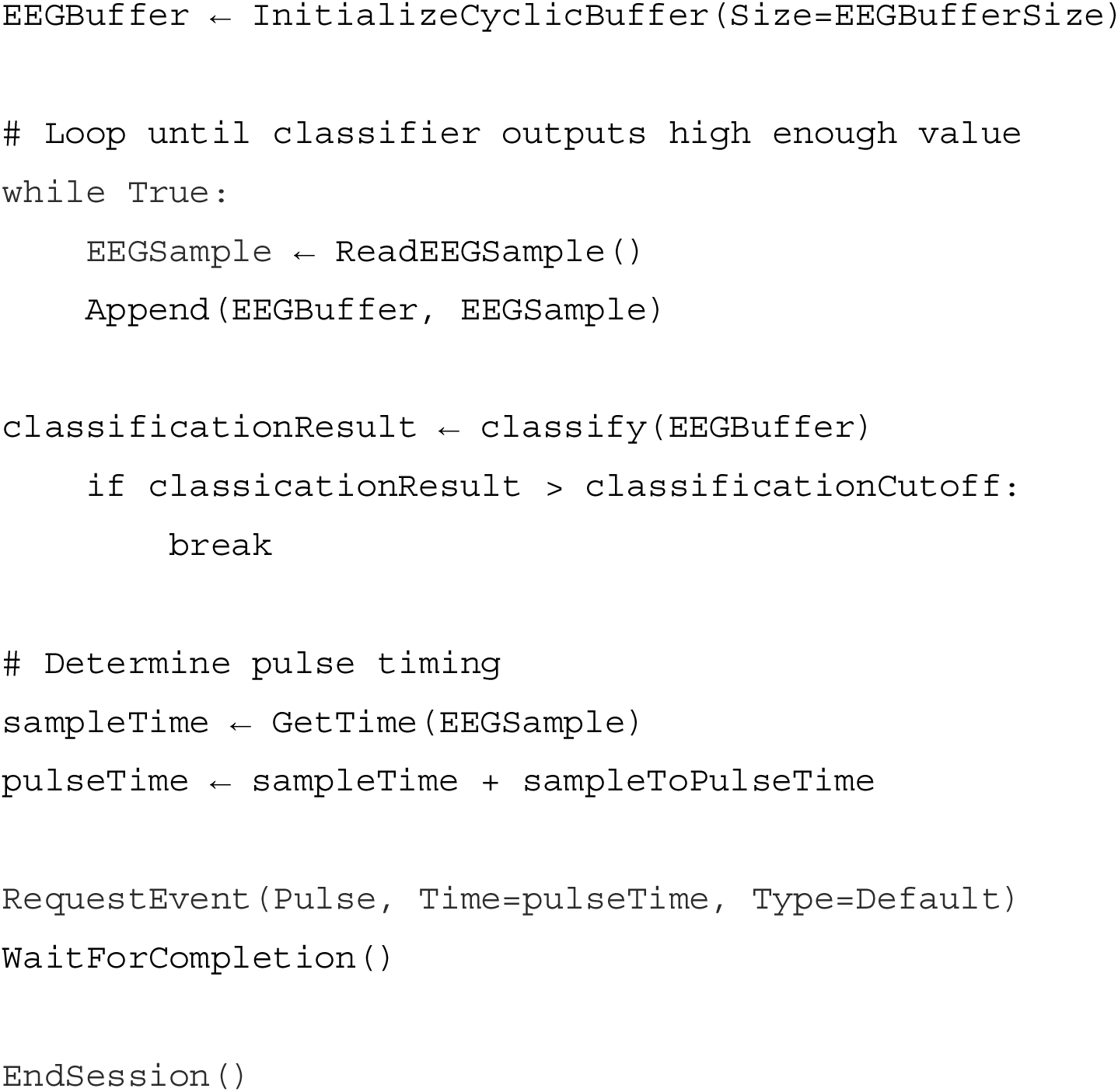

## Footnotes

1 Magstim BiStim

2 https://www.magstim.com/product/magstim-bistim2/

## References

[1] Barker A T, Jalinous R and Freeston I L 1985 Non-invasive magnetic stimulation of human motor cortex The Lancet 325 1106–7

[2] Ilmoniemi R J, Ruohonen J and Karhu J 1999 Transcranial magnetic stimulation–a new tool for functional imaging of the brain Crit Rev Biomed Eng 27 241–84

[3] Vucic S, Stanley Chen K H, Kiernan M C, Hallett M, Benninger D H, Di Lazzaro V, Rossini P M, Benussi A, Berardelli A, Currà A, Krieg S M, Lefaucheur J P, Long Lo Y, Macdonell R A, Massimini M, Rosanova M, Picht T, Stinear C M, Paulus W, Ugawa Y, Ziemann U and Chen R 2023 Clinical diagnostic utility of transcranial magnetic stimulation in neurological disorders. Updated report of an IFCN committee Clinical Neurophysiology 150 131–75

[4] Lefaucheur J P, Aleman A, Baeken C, Benninger D H, Brunelin J, Di Lazzaro V, Filipović S R, Grefkes C, Hasan A, Hummel F C, Jääskeläinen S K, Langguth B, Leocani L, Londero A, Nardone R, Nguyen J P, Nyffeler T, Oliveira-Maia A J, Oliviero A, Padberg F, Palm U, Paulus W, Poulet E, Quartarone A, Rachid F, Rektorová I, Rossi S, Sahlsten H, Schecklmann M, Szekely D and Ziemann U 2020 Evidence-based guidelines on the therapeutic use of repetitive transcranial magnetic stimulation (rTMS): an update (2014–2018) Clinical Neurophysiology 131 2150–206

[5] Kundu B, Johnson J S and Postle B R 2014 Prestimulation phase predicts the TMS-evoked response J Neurophysiol 112 1885–93

[6] Kraus D, Naros G, Bauer R, Khademi F, Leão M T, Ziemann U and Gharabaghi A 2016 Brain state-dependent transcranial magnetic closed-loop stimulation controlled by sensorimotor desynchronization induces robust increase of corticospinal excitability Brain Stimul 9 415–24

[7] Zrenner C, Belardinelli P, Müller-Dahlhaus F and Ziemann U 2016 Closed-loop neuroscience and non-invasive brain stimulation: a tale of two loops Front Cell Neurosci 10 92

[8] Rösch J, Emanuel Vetter D, Baldassarre A, Souza V H, Lioumis P, Roine T, Jooß A, Baur D, Kozák G, Blair Jovellar D, Vaalto S, Romani G L, Ilmoniemi R J and Ziemann U 2024 Individualized treatment of motor stroke: a perspective on open-loop, closed-loop and adaptive closed-loop brain state-dependent TMS Clinical Neurophysiology 158 204–11

[9] Zrenner C, Desideri D, Belardinelli P and Ziemann U 2018 Real-time EEG-defined excitability states determine efficacy of TMS-induced plasticity in human motor cortex Brain Stimul 11 374–89

[10] Humaidan D, Xu J, Kirchhoff M, Romani G L, Ilmoniemi R J and Ziemann U 2024 Towards real-time EEG–TMS modulation of brain state in a closed-loop approach Clinical Neurophysiology 158 212–7

[11] Bergmann T O 2018 Brain state-dependent brain stimulation Front Psychol 9 2108

[12] Marzetti L, Makkinayeri S, Pieramico G, Guidotti R, D’Andrea A, Roine T, Mutanen T P, Souza V H, Kičić D, Baldassarre A, Ermolova M, Pankka H, Ilmoniemi R J, Ziemann U, Luca Romani G and Pizzella V 2024 Towards real-time identification of large-scale brain states for improved brain state-dependent stimulation Clinical Neurophysiology 158 196–203

[13] Pineda J A 2005 The functional significance of mu rhythms: translating “seeing” and “hearing” into “doing” Brain Res Rev 50 57–68

[14] Stefanou M I, Baur D, Belardinelli P, Bergmann T O, Blum C, Gordon P C, Nieminen J O, Zrenner B, Ziemann U and Zrenner C 2019 Brain state-dependent brain stimulation with real-time electroencephalography-triggered transcranial magnetic stimulation Journal of Visualized Experiments 2019 150

[15] Varone G, Boulila W, Pascarella A, Gasparini S and Aguglia U 2024 Instrumentation for TMS-EEG Experiment: ArTGen and a custom EEG interface Int J Imaging Syst Technol 34 e23134

[16] Barry J R, Lee E A and Messerschmitt D G 2012 Digital Communication (Springer)

[17] Scheer G W and Moxley R E 2006 Digital communications improve contact I/O reliability Proceedings of the Power Systems Conference: Advanced Metering, Protection, Control, Communication, and Distributed Resources pp 363–77

[18] Viterbi A J and Omura J K 2009 Principles of Digital Communication and Coding (Courier Corporation)

[19] Ben Mahmoud M S, Pirovano A and Larrieu N 2014 Aeronautical communication transition from analog to digital data: a network security survey Comput Sci Rev 11 1–29

[20] Angerer A, Hoffmann A, Schierl A, Vistein M and Reif W 2013 Robotics API: object-oriented software development for industrial robots Journal of Software Engineering for Robotics 4 1–22

[21] Universal Robots 2015 The URScript Programming Language (Technical Manual)

[22] Moog R A 1986 MIDI: Musical Instrument Digital Interface Journal of the Audio Engineering Society 34 394–404

[23] Loy G 1985 Musicians make a standard: the MIDI phenomenon Computer Music Journal 9 8–26

[24] Hsu J Y 2002 Transistor circuits In Computer Logic: Design Principles and Applications (Springer) pp 62–90

[25] Deep A, Kim J and Shyam S K 2014 Optimized USB 2.0 scheduler for low latency data transfer Proceedings of the International Symposium on Consumer Electronics (ISCE) pp 1–2

[26] Horowitz P, Hill W and Robinson I 1989 The Art of Electronics vol 2 (Cambridge University Press)

[27] Roscoe A 1998 The Theory and Practice of Concurrency (Prentice Hall)

[28] Hassan U, Pillen S, Zrenner C and Bergmann T O 2022 The Brain Electrophysiological recording & STimulation (BEST) toolbox Brain Stimul 15 109–15

[29] Ramadoss L and Hung J Y 2008 A study on Universal Serial Bus latency in a real-time control system Proceedings of the Industrial Electronics Conference (IECON) pp 67–72

[30] Piessens F, Verhanneman T and Win B De 2002 On the importance of the separation-of-concerns principle in secure software engineering Proceedings of the Workshop on the Application of Engineering Principles to System Security Design 1–10

[31] Habibollahi Saatlou F, Rogasch N C, McNair N A, Biabani M, Pillen S D, Marshall T R and Bergmann T O 2018 MAGIC: An open-source MATLAB toolbox for external control of transcranial magnetic stimulation devices Brain Stimul 11 1189–91

[32] McNair N A 2017 MagPy: a Python toolbox for controlling Magstim transcranial magnetic stimulators J Neurosci Methods 276 33–7

[33] Teng F C 2000 Real-time control using MATLAB Simulink Proceedings of the IEEE International Conference on Systems, Man and Cybernetics vol 4 pp 2697–702

[34] Pankka H, Lehtinen J, Ilmoniemi R J and Roine T 2024 Enhanced EEG forecasting: a probabilistic deep learning approach [preprint] bioRxiv 2024.01.16.575836

[35] Stenroos M and Koponen L M 2019 Real-time computation of the TMS-induced electric field in a realistic head model Neuroimage 203 116159

[36] Kida T, Tanaka E and Kakigi R 2016 Multi-dimensional dynamics of human electromagnetic brain activity Front Hum Neurosci 9 713

[37] Reghenzani F, Massari G and Fornaciari W 2019 The real-time Linux kernel: a survey on PREEMPT_RT ACM Comput Surv 52 1–36

[38] Kahilakoski O-P, Alkio K, Siljamo O, Valén K, Laurinoja J, Haxel L, Makkonen M, Mutanen T P, Tommila T, Guidotti R, Pieramico G, Ilmoniemi R J and Roine T 2024 NeuroSimo: an open-source software for closed-loop EEG- or EMG-guided TMS [submitted] Brain Stimul

[39] Lieb A, Zrenner B, Zrenner C, Kozák G, Martus P, Grefkes C and Ziemann U 2023 Brain-oscillation-synchronized stimulation to enhance motor recovery in early subacute stroke: a randomized controlled double-blind three-arm parallel-group exploratory trial comparing personalized, non-personalized and sham repetitive transcranial magnetic stimulation (acronym: BOSS-STROKE) BMC Neurol 23 204

[40] Bovy C J, Mertodimedjo H T, Hooghiemstra G, Uijterwaal H and Van Midghem P 2002 Analysis of end-to-end delay measurements in Internet Proceedings of the ACM Conference on Passive and Active Measurements (PAM)

[41] Shore J 2004 Fail fast [software debugging] IEEE Softw 21 21–5

[42] Martins L E G and Gorschek T 2022 Requirements Engineering for Safety-Critical Systems (CRC Press)

[43] Koopman P and Wagner M 2014 Transportation CPS safety challenges Proceedings of the NSF Workshop on Transportation CyberPhysical Systems (Carnegie Mellon University)

[44] Laplante P A and Ovaska S J 2011 Real-time Systems Design and Analysis (Wiley)

[45] Santhosh R and Ravichandran T 2013 Pre-emptive scheduling of on-line real time services with task migration for cloud computing Proceedings of the International Conference on Pattern Recognition, Informatics and Mobile Engineering (PRIME) pp 271–6

[46] MIDI Manufacturers Association Summary of MIDI 1.0 messages (Technical Specification)

[47] MIDI Manufacturers Association Standard MIDI-file format specification 1.1 (Technical Specification)

[48] Tanenbaum A S and Van Steen M 2017 Distributed Systems (CreateSpace)

[49] Mills D L 1991 Internet time synchronization: the Network Time Protocol IEEE Transactions on Communications 39 1482–93

[50] Girela-López F, López-Jiménez J, Jiménez-López M, Rodríguez R, Ros E and Díaz J 2020 IEEE 1588 High Accuracy Default Profile: applications and challenges IEEE Access 8 45211–20

[51] Watt S T, Achanta S, Abubakari H, Sagen E, Korkmaz Z and Ahmed H 2016 Understanding and applying precision time protocol Proceedings of the Saudi Arabia Smart Grid (SASG) pp 1–7

[52] Horauer M, Kerö N and Schmid U 2000 A Network Interface for Highly Accurate Clock Synchronization (Technical Report)

[53] Nieminen J O, Sinisalo H, Souza V H, Malmi M, Yuryev M, Tervo A E, Stenroos M, Milardovich D, Korhonen J T, Koponen L M and Ilmoniemi R J 2022 Multi-locus transcranial magnetic stimulation system for electronically targeted brain stimulation Brain Stimul 15 116–24

[54] Marouani H and Dagenais M R 2008 Internal clock drift estimation in computer clusters Journal of Computer Networks and Communications 2008 583162

[55] Bittium Oyj 2024 NeurOne System User Manual (User Manual)

[56] Koponen L M, Nieminen J O and Ilmoniemi R J 2018 Multi-locus transcranial magnetic stimulation—theory and implementation Brain Stimul 11 849–55

[57] Navarro de Lara L I, Daneshzand M, Mascarenas A, Paulson D, Pratt K, Okada Y, Raij T, Makarov S N and Nummenmaa A 2021 A 3-axis coil design for multichannel TMS arrays Neuroimage 224 117355

[58] Sinisalo H, Kahilakoski O-P, Souza V H, Nieminen J O, Rantala R, Tommila T, Usuga I, Laine M, Ahola O, Gallegos E, Kozák G, Vetter D E, Rissanen I, Jooß A, Matsuda R, Soto A M, Li D, Humaidan D, Stenroos M, Roine T, Kičić D, Ziemann U and Ilmoniemi R J 2024 Design, construction, and deployment of a multi-locus transcranial magnetic stimulation system for clinical use [submitted] IEEE Trans Biomed Eng

[59] Zhu X, Song B, Ni Y, Ren Y and Li R 2016 Software defined anything—from software-defined hardware to software defined anything Business Trends in the Digital Era pp 83–103

[60] Kreutz D, Ramos F M V, Verissimo P E, Rothenberg C E, Azodolmolky S and Uhlig S 2015 Software-defined networking: a comprehensive survey Proceedings of the IEEE 103 14–76

[61] Mei H and Guo Y 2018 Toward ubiquitous operating systems: A software-defined perspective Computer (Long Beach Calif) 51 50–6

[62] De Michell G and Gupta R K 1997 Hardware/software co-design Proceedings of the IEEE 85 349–65

[63] Wolf W H 1994 Hardware-software co-design of embedded systems Proceedings of the IEEE 82 967–89

[64] Hernandez-Pavon J C, Veniero D, Bergmann T O, Belardinelli P, Bortoletto M, Casarotto S, Casula E P, Farzan F, Fecchio M, Julkunen P, Kallioniemi E, Lioumis P, Metsomaa J, Miniussi C, Mutanen T P, Rocchi L, Rogasch N C, Shafi M M, Siebner H R, Thut G, Zrenner C, Ziemann U and Ilmoniemi R J 2023 TMS combined with EEG: recommendations and open issues for data collection and analysis Brain Stimul 16 567–93

[65] Matsuda R H, Souza V H, Marchetti T C, Soto A M, Kahilakoski O-P, Zhdanov A, Malheiro V H E, Laine M, Nyrhinen M, Sinisalo H, Kicic D, Lioumis P, Ilmoniemi R J and Baffa O 2024 Robotic–electronic platform for autonomous and accurate transcranial magnetic stimulation targeting Brain Stimul 17 469–72

[66] Memarian Sorkhabi M, Wendt K, O’Shea J and Denison T 2022 Pulse width modulation-based TMS: primary motor cortex responses compared to conventional monophasic stimuli Brain Stimul 15 980–3

[67] Sinisalo H, Laine M, Nieminen J O, Souza V H, Matsuda R H, Soto A M, Ukharova E, Mutanen T, Rissanen I, Stenroos M, Koponen L M and Ilmoniemi R J 2024 Multi-locus transcranial magnetic stimulation with pulse-width modulation [submitted] Brain Stimul

